# Estrogen receptor activation remodels *TEAD1* gene expression to alleviate nonalcoholic fatty liver disease

**DOI:** 10.1101/2023.09.07.556687

**Authors:** Christian Sommerauer, Carlos J. Gallardo-Dodd, Christina Savva, Linnea Hases, Madeleine Birgersson, Rajitha Indukuri, Joanne X. Shen, Pablo Carravilla, Keyi Geng, Jonas Nørskov Søndergaard, Clàudia Ferrer-Aumatell, Grégoire Mercier, Erdinc Sezgin, Marion Korach-André, Carl Petersson, Hannes Hagström, Volker M. Lauschke, Amena Archer, Cecilia Williams, Claudia Kutter

## Abstract

**Introduction:** The occurrence of obesity-related hepatic malignancies differs between sexes, suggesting the involvement of sex hormones. Female sex hormones maintain cell homeostasis through estrogen receptor (ER) signaling and protect from developing nonalcoholic fatty liver disease (NAFLD) in mice and humans.

**Rationale:** To understand recovery from high-fat diet (HFD)-induced liver disease in males upon estrogen treatment, we comprehensively characterized molecular changes in the liver upon selective activation of estrogen receptors (ERs) to identify novel therapeutic targets downstream of estrogen signaling.

**Methods:** To dissect hepatic ER isoform-driven responses, we integrated liver transcriptomes from female and male HFD mice treated with or without four different estrogen agonists, along with multiomics data, including bulk, single-cell and spatial transcriptomics, chromatin profiling, machine learning models and advanced microscopy. Patient cohorts and primary human hepatocyte spheroids datasets were included.

**Results:** Only males developed liver steatosis. We found that selective activation of either ERα or ERβ reduced HFD-induced hepatic steatosis in male mice. Systemic ER activation restored HFD-induced aberrant gene expression of cellular processes across liver cell types, including hepatocytes. Profiling of marked histones revealed that ER activation modulated promoter and enhancer sites and identified 68 estrogen-sensitive enhancer-gene pairs. Most of these genes were similarly deregulated in human nonalcoholic fatty liver disease (NAFLD) patients, including the transcription factor *TEAD1*. *TEAD1* expression increased in NAFLD patients, and inhibiting TEAD ameliorated steatosis in spheroids by suppressing lipogenic pathways.

**Conclusions:** Systemic activation of ERα or ERβ modulates molecular pathways in the liver to counteract NAFLD. Our study identified *TEAD1* as a key ER-sensitive gene and suggested that its inhibition poses a therapeutic strategy to combat NAFLD without the undesired side effects elicited by estrogen signaling.

**Clinical research relevance:** We identified drug targets downstream of estrogen signaling, including TEAD1, and demonstrate that TEAD inhibition improves steatosis by suppressing lipogenic pathways.

**Basic research relevance:** The targeted activation of nuclear ERs recovers high-fat diet-induced molecular and physiological liver phenotypes by remodeling core pathways beyond lipid metabolism. ER-responsive enhancers regulate central metabolic genes of clinical significance in NAFLD patients, highlighting the potential impact of this research on understanding liver cell plasticity.

**HIGHLIGHTS:**

- steatosis in livers of high-fat diet (HFD) male mice was effectively reduced by selective activation of estrogen receptors (ERα and ERβ) with four different agonists.
- ER agonist treatments successfully reversed HFD-induced changes in gene regulation and expression, revealing new treatment targets involving previously unconnected molecular pathways.
- estrogen-sensitive enhancers regulated important genes, including TEAD1, emerging as pivotal NAFLD regulators significantly impacting metabolic processes.
- high *TEAD1* gene expression in NAFLD patients correlated with disease severity, underscoring its clinical significance in disease progression.
- inhibiting TEAD with small molecules alleviated steatosis by suppressing lipogenic pathways, resembling some of the same beneficial effects as estrogen treatment.

## BACKGROUND

The global obesity epidemic poses a substantial risk for metabolic disorders, including liver diseases [1]. Prolonged high-calorie diets, like high-fat diet (HFD), induce hepatic lipid accumulation, resulting in hepatic steatosis, the defining hallmark of nonalcoholic fatty liver disease (NAFLD). Persistent dietary imbalance causes steatohepatitis (NASH), characterized by hepatocyte death, inflammation, and progressive liver fibrosis, potentially developing into cirrhosis and liver cancer [2]. NAFLD prevalence has risen alongside obesity, currently affecting one-third of adults worldwide [1]. Yet, approved medications for NAFLD treatment are lacking, highlighting the urgency to identify suitable targets.

NAFLD occurrence differs greatly between sexes, with lower prevalence in premenopausal women than in men or postmenopausal women [3]. The female sex hormone estrogen exerts protective roles in the liver, but the underlying molecular mechanisms remain understudied [4]. Estrogens bind to nuclear estrogen receptors (ERα and ERβ), acting as transcription factors that activate or repress target genes and signaling cascades by either direct DNA interaction or tethering to other transcription factors [4,5].

Estrogen signaling is crucial in females and males. Endogenous estrogen is produced by enzymatic cholesterol conversion in both sexes. In male mice on a conventional diet, the deficiency of the enzyme aromatase leads to hepatic steatosis [6], and similarly, liver-specific ERα impairment also induces abnormal liver physiology and liver energy metabolism [7,8]. Menopausal hormone therapy in women reduces NAFLD prevalence, highlighting that estrogen signaling safeguards hepatic energy metabolism [3]. Modulating estrogen levels or ER activity affects hepatic molecular changes and NAFLD susceptibility [9]. Identifying estrogen-responsive factors and pathways can enhance treatment options for obesity-related liver morbidities while avoiding potential estrogen treatment side effects [10].

In this study, we identified sex-specific molecular signatures that link the hepatoprotective role of ERs to downstream effectors in a diet-induced NAFLD mouse model. Utilizing an integrative multiomics approach, we examined transcriptional and chromatin changes in liver leveraging on single-cell and spatial information. Systemic activation of ER isoforms in mice elucidated their distinct hepatoprotective effects. We found that ER-controlled murine key factors, including *TEAD1*, were similarly altered in NAFLD patients. We demonstrated that small molecule-based TEAD inhibition reduced lipid accumulation in an organotypic human liver model by suppressing lipogenesis. Collectively, we identified gene regulatory circuits downstream of ER-signaling that control hepatic metabolism and determined that network signature-informed interference can ameliorate liver disease phenotypes.

## RESULTS

### HFD severely changed molecular and physiological parameters in male C57BL/6J mice

To assess diet-induced sexual dimorphism in liver transcriptomes resembling early NAFLD stages, we fed five-week-old female and male C57BL/6J inbred mice a control (CD, 10% fat) or high-fat diet (HFD, 60% fat) for 13 weeks (**Fig. 1A**). Upon HFD, both sexes gained weight [11]. Males but not females on HFD developed hepatic steatosis, increased liver weight and circulating glucose levels (**Fig. 1B**, **Supplemental Fig. 1, A-D** and **Supplemental Table 1**). These findings confirmed that female mice were more protected from HFD than males.

**Fig. 1.**
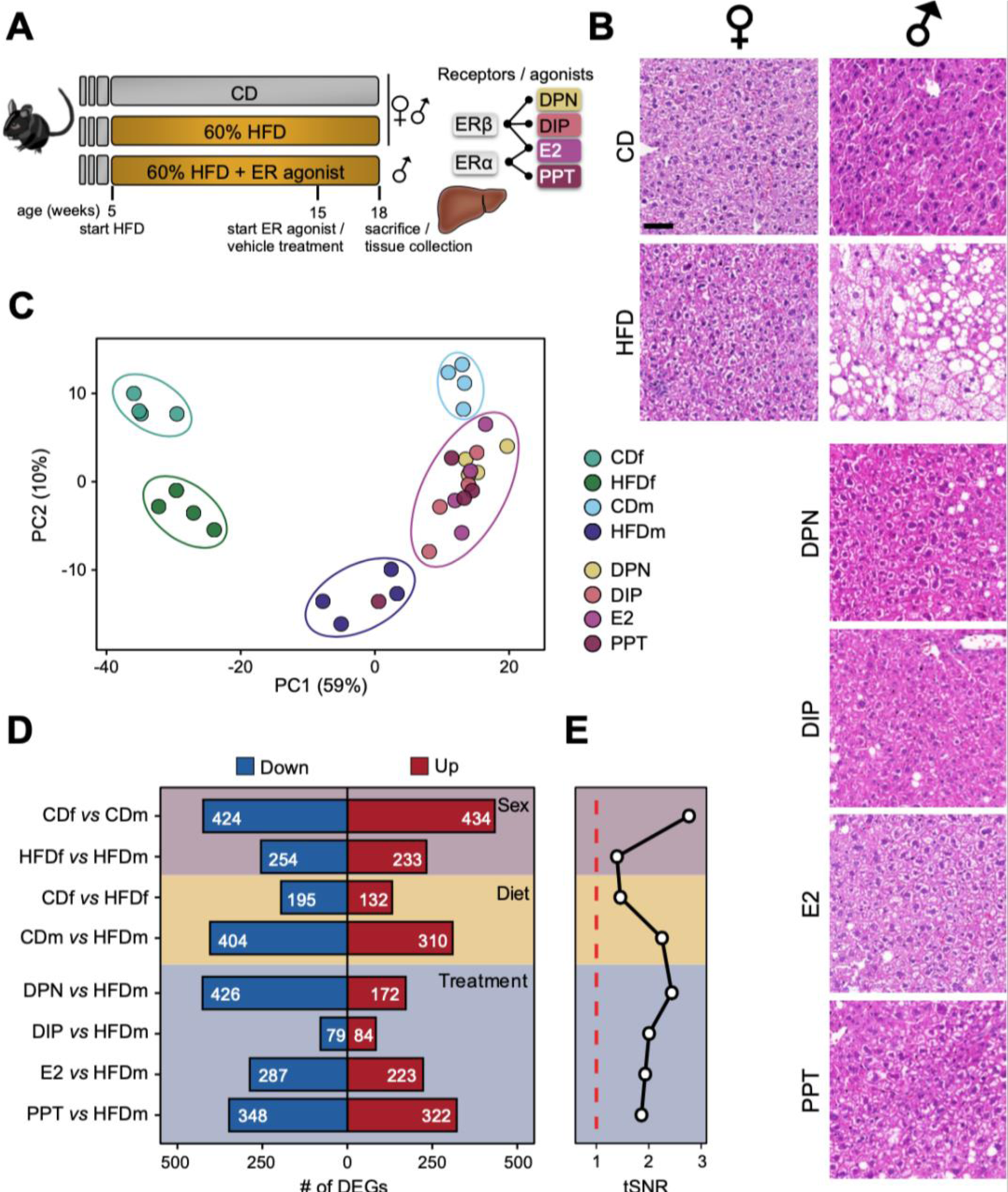
Male mice are severely affected by high-fat diet. **(A)** Schematic representation of the mouse experimentation. Five-week-old female (f) and male (m) C57BL/6 mice received either a control (CD, 10% fat) or high-fat diet (HFD, 60% fat) for 13 weeks. HFDm subgroups were injected with estrogen receptor α (ERα, E2 or PPT) or ERβ (DPN or DIP) agonists every other day from week 15 to 18. Isolated livers were histologically and molecularly assessed. **(B)** Liver cross-sections of female (left) and male (right) mice on different diets and ER-agonist treatments were stained with hematoxylin and eosin. Scale bar: 50μm. **(C)** Factorial map of the principal components (PC) analysis separates global gene expression levels. Percentage of PC variance is shown (parentheses). Color-coded small circles illustrate individual mice on different diets and treatments. Color-coded large ellipses group mice by sex, diet, and treatment. **(D)** Horizontal bars present the number (highlighted) of downregulated (blue) and upregulated (red) genes for sex (purple), diet (yellow) or treatment (blue) comparisons. **(E)** Black line shows transcriptome-based signal-to-noise ratio (tSNR, x-axis) for Fig. 1d comparisons. Dashed red line represents the noise baseline.

To investigate the underlying molecular effects, we profiled the transcriptome of livers from male and female mice on CD and HFD (n=4) (**Supplemental Tables 2-4**). Our principal component analysis (PCA) separated our samples primarily by sex (PC1, 59%) and by diet (PC2, 10%) (**Fig. 1C**). HFD males exhibited more differentially expressed genes (DEGs) (n=714) than HFD females (n=327) demonstrating that gene expression in males was more susceptible to HFD than in females, irrespective of genes expressed on the sex chromosomes (**Fig. 1D, Diet, Supplemental Table 2**). Only a fraction of genes was commonly deregulated between females and males on HFD, further emphasizing the sex disparity in response to dietary stimuli (**Supplemental Fig. 1E**).. We further confirmed these findings by quantifying threshold-independent differences for each comparison (**Fig. 1E, Diet**). Genes deregulated in both sexes or in HFD males were enriched in biological processes (gene ontology, GO) linked to lipid metabolism, while HFD females exhibited enrichment in circadian rhythm (**Supplemental Fig. 1E** and **Supplemental Table 5**).

Taken together, we found that male mice responded stronger to HFD than females and these differences could be traced back to major sex differences in liver transcriptomes.

### Systemic ER activation mitigates diet-induced liver alterations in an isoform-specific manner

Given the resilience of female mice to HFD, we tested the hepatoprotective effects of estrogen in males. After ten weeks, HFD male mice were injected with agonists that selectively activate ERβ (DPN and DIP) [40,41], ERα (PPT) [40] or both (E2) [40] every other day for three weeks (**Fig. 1A**). As previously reported, estrogenic ligand treatments modestly reduced total weight, liver weight and blood glucose levels (**Supplemental Fig. 1, A-C**) [11]. Strikingly, all ER-agonists reduced steatosis compared to vehicle-treated HFD males (**Fig. 1B** and **Supplemental Fig. 1D**). Our PCA showed that agonist-treated HFD males clustered between HFD and CD males, implying attenuation of HFD-induced alterations (**Fig. 1C**). DIP had the weakest impact on the transcriptome (n=163 DEGs), whereas DPN (n=598 DEGs), E2 (n=510 DEGs) and PPT (n=670 DEGs) had greater effects (**Fig. 1, D** and **E**). DPN predominantly downregulated genes, while DIP, E2, and PPT treatments had similar proportions of down- and upregulated genes (**Fig. 1D**).

We unified DEGs across all male comparisons (n=1,477), which formed four expression clusters (**Fig. 2A**). Cluster 1 (n=577) exhibited HFD-induced gene upregulation compared to CD, attenuated by all agonists, while cluster 2 (n=258) displayed HFD-induced gene upregulation, partially restored upon agonist treatment. Cluster 3 (n=295) contained genes with higher HFD expression and ERβ-dependent repression, and cluster 4 (n=346) included genes upregulated by ERα.

**Fig. 2.**
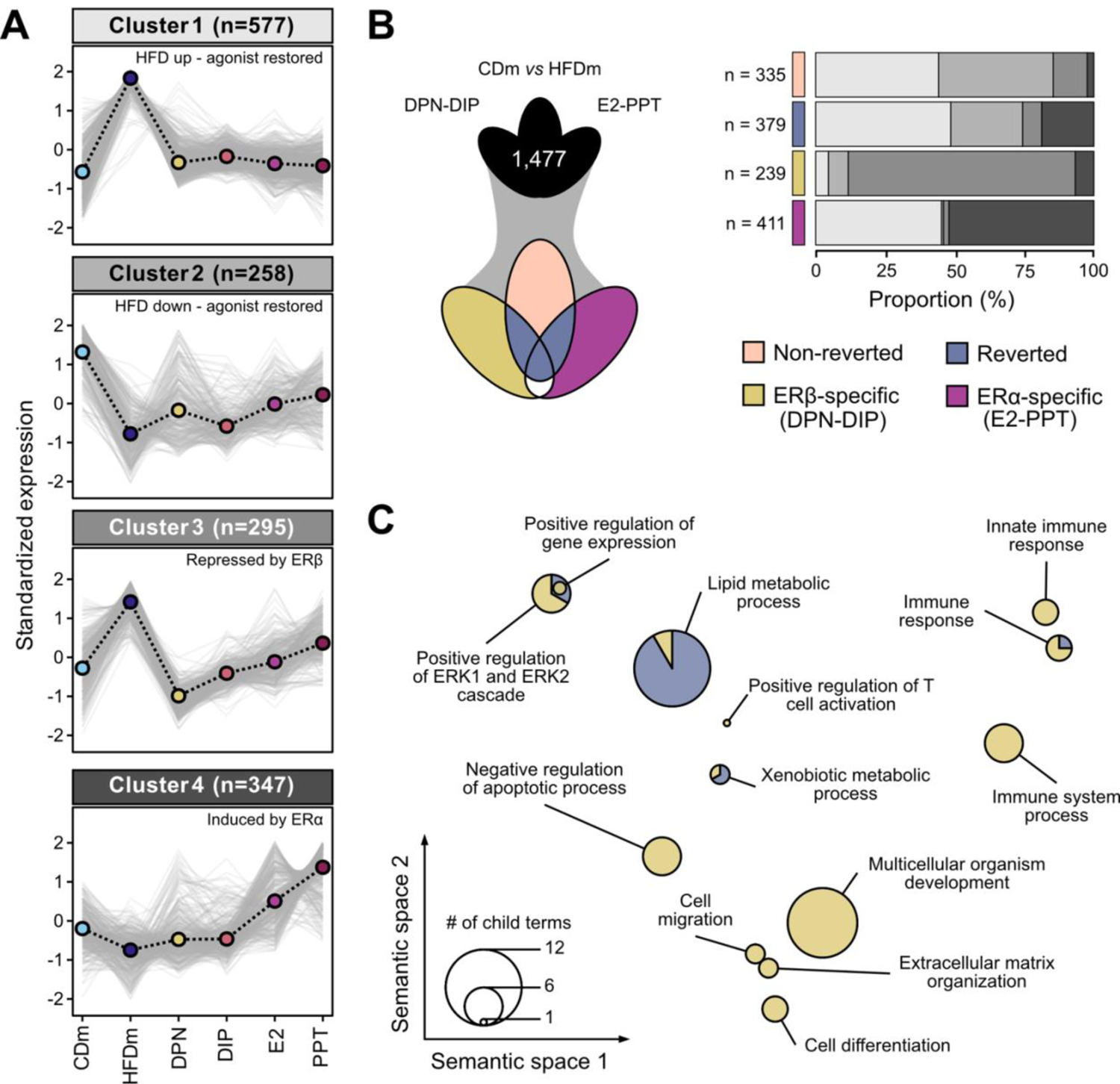
ERα/β-agonist treatment largely reverts HFD-induced transcriptome alterations in males. **(A)** Line charts depict four clusters (grey-scaled) of gene expression trends (*z*-score) for unified deregulated genes (DEGs, n=1,477) in mice on different diets and ER-agonist treatments (color-coded). Cluster centroid (dashed black line) represents all deregulated genes (grey lines). Number of genes per cluster is shown (parentheses). **(B)** Three-way Venn diagram (left) presents intersections of unified DEGs (clustered in Fig. 2a). Genes are categorized (right) into non-reverted (rose), reverted (denim), ERβ-specific (DPN-DIP, ochre) and ERα-specific (E2-PPT, violet) gene sets. Horizontal bar chart displays the proportional occurrences of gene sets in the four clusters. Number (n) indicates gene set size. **(C)** PCA factorial map separates the semantic space of enriched gene ontology (GO) terms in reverted and ERβ-specific gene sets (circles). Enriched GO terms are collapsed at the parent term level and separated based on similarity (x-y-axis). Circle size corresponds to number of enriched GO terms.

We next stratified the DEGs into four categories (**Fig. 2B** and **Supplemental Table 6**). Genes significantly deregulated by HFD were termed ‘non-reverted’ (n=333) when unaffected by ER-agonist treatment and ‘reverted’ (n=381) when restored by at least one treatment. Most of these genes resided in clusters 1 and 2, suggesting an overall adjustment towards the CD state (**Fig. 2B**). Additionally, we distinguished ‘ERβ-specific’ (DPN-DIP, n=239) and ‘ERα-specific’ (E2-PPT, n=411) gene signatures with unchanged expression levels upon HFD but altered upon ER-agonist treatment. Although E2 activates ERα and ERβ, we found a higher overlap between E2- and PPT-than E2- and DPN-regulated genes indicating that E2 primarily acted through ERα (**Supplemental Fig. 2, A** and **B**). ERβ-specific genes were mostly in cluster 3, while ERα-specific genes were predominantly in clusters 1 and 4 (**Fig. 2B**). The degree of recovery varied among ER-agonist treatments, with PPT and DPN showing the highest number of reversed HFD-deregulated genes (38% and 37%, respectively), followed by E2 (35%) and DIP (16%) (**Supplemental Fig. 2C**).

For each of the four categories, we investigated gene enrichments in GO biological processes. The reverted and ERβ-specific gene sets showed significant enrichments of genes regulating lipid metabolism, ERK signaling, xenobiotic metabolism and immune responses (**Fig. 2C, Supplemental Fig. 2D** and **Supplemental Table 5**). Additionally, the ERβ-specific gene sets controlled extracellular matrix organization, apoptosis, cell motility and differentiation processes, which were almost entirely represented in cluster 3 characterized by ERβ-agonist treatment-specific gene downregulation (**Fig. 2B** and **Supplemental Fig. 2D**). We found no GO term overrepresentation for non-reverted and ERα-specific gene categories.

Overall, systemic ERα or ERβ activation restored diet-induced gene expression changes, with isoform-specific differences, correcting metabolic processes to reduce steatotic phenotypes (**Fig. 1B**).

### Systemic ER activation has widespread implications in core liver pathways

We performed a threshold-independent gene set enrichment analysis (GSEA) to capture functionally relevant genes recovered upon ER-agonist treatments but without reaching statistical significance (**Fig. 2**). Reactome pathway analysis, clustering, and subsequent correlation based on normalized enrichment scores (NES), identified 24 relevant pathway clusters that were significantly altered in HFD males compared to CD males and HFD ER-agonist-treated males (**Fig. 3A, Supplemental Fig. 3** and **4**, and **Supplemental Table 7**).

**Fig. 3.**
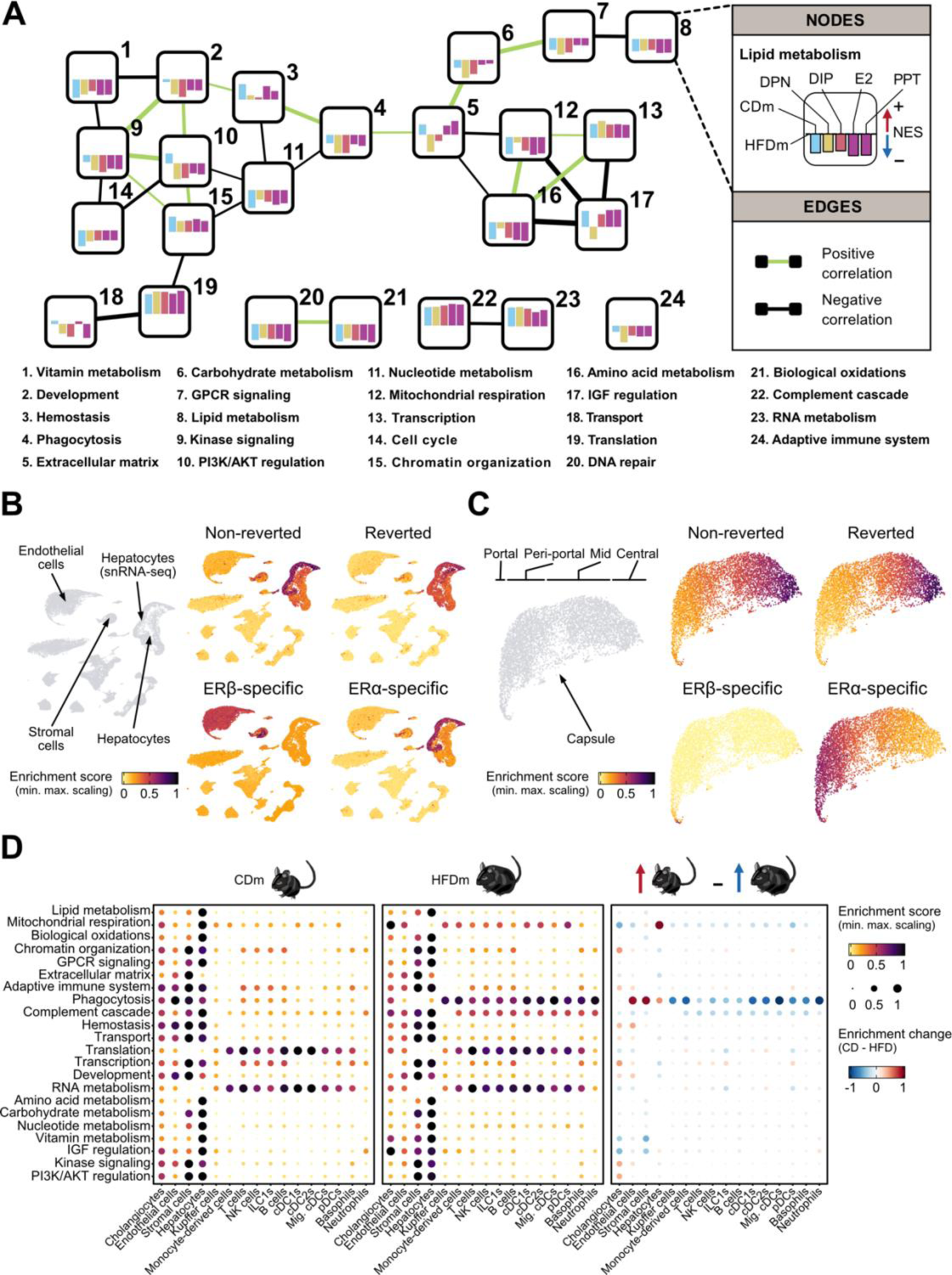
ERα/β-agonist treatment reverts HFD-induced transcriptional changes by affecting central cellular pathways and liver cell types. (**A**) Network connects major Reactome pathway clusters (numbered nodes). Colored bars inside each node present pathway cluster enrichment for CDm and ER-agonist-treated HFDm compared to HFDm (normalized enrichment score, NES). Edges connect positively-correlated (green) or negatively-correlated (black) nodes based on NES profiles. (**B**) UMAP space-projected enrichment plots highlight liver cell types with enhanced signal for the gene sets (defined in Fig. 2b). Hepatocyte nuclear fraction (snRNA-seq) is labelled. (**C**) Spatial transcriptomics maps show liver zonation patters of the gene sets (Fig. 2b). (**D**) Bubble plots display activity scores of altered pathway clusters (Fig. 3a) across all liver cell types in control (left), HFD (middle) male mice and their differences (right). Color-code and circle diameter for enrichment score: low (yellow, narrow), high (black, wide). Arrows and enrichment change indicate higher abundance in control (red) and HFD (blue) mice.

**Fig. 4.**
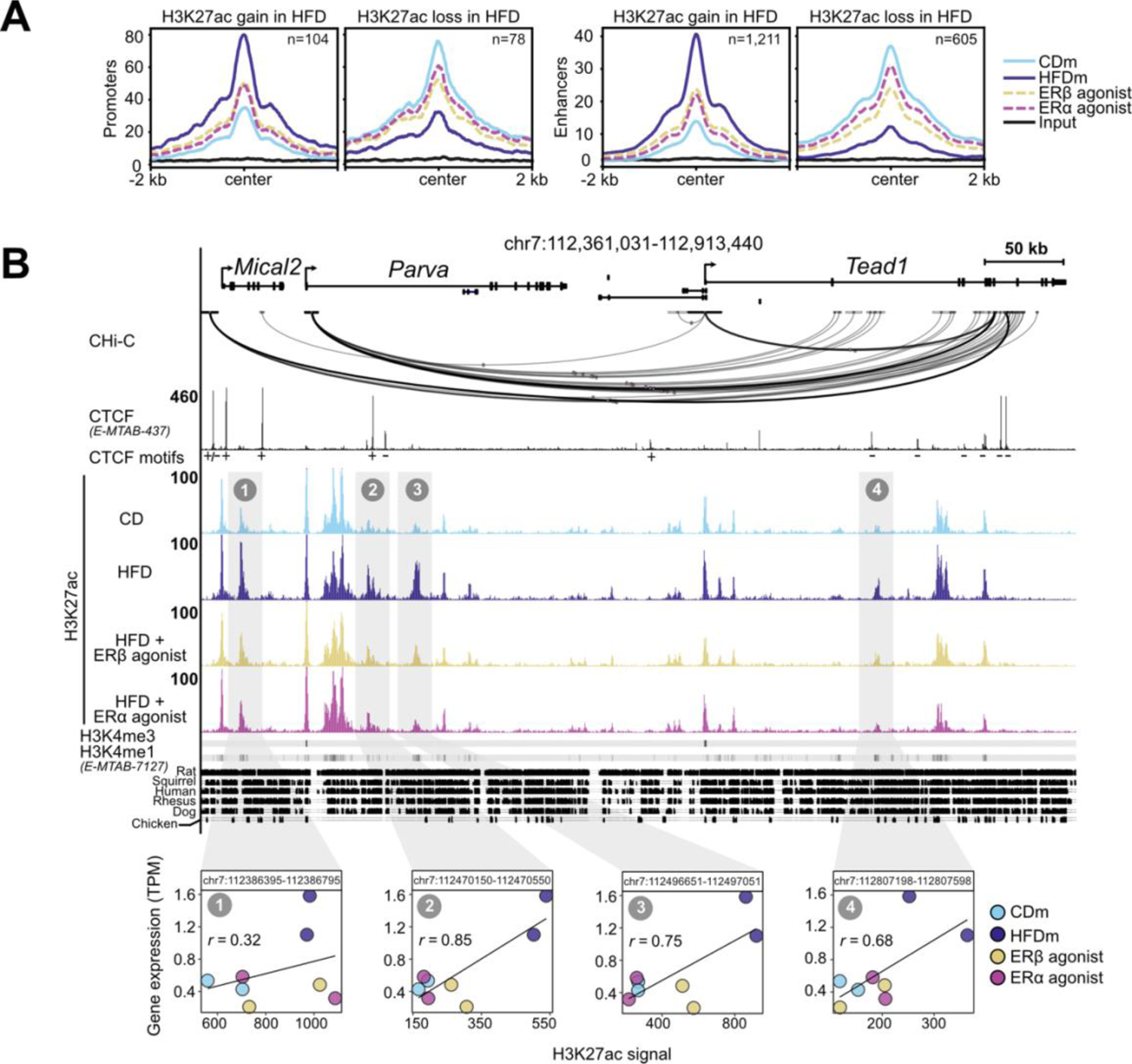
ERα/β-agonist treatment recovers HFD-induced changes at enhancers and promoters. **(A)** Metaplots show H3K27ac read aggregation in promoters (left) and enhancers (right), centered at the peak summits. Number (n) indicates significant H3K27ac signal gains or losses in livers of HFDm compared to CDm. One representative replicate per condition is depicted. **(B)** Genome browser view (mm10) illustrates genomic region around the *Tead1* gene locus. Black boxes represent exons and UTRs. Arrows indicated gene transcription directionality. Scale bar shows genomic region length in kilobases (kb). Black arcs display promoter-capture HiC (C-HiC) 3D connections. Genomic regions are enriched for CTCF (black peaks) with CTCF motif orientations (plus or minus symbols), H3K27ac (color-coded peaks) in CDm, HFDm and HFDm treated with ERβ-agonist (DPN) or ERα-agonist (E2), H3K4me3 and H3K4me1 (horizonal grey bar, dark: high, light: low). The y-axis of each track specifies normalized read density. Enhancer locations paired with the *Tead1* gene locus are highlighted (grey vertical boxes). The degree of genomic sequence conservation in vertebrates is shown (conserved: black, not conserved: white). Scatter plots correlate *Tead1* gene expression (TPM, y-axis) and its paired enhancers (H3K27ac signal, x-axis) in livers of male mice on different diets and ER-agonist treatments. All replicates are shown. Enhancer coordinates (400 bp window around the enhancer summit) and Pearson correlation coefficients (*r*) are indicated in each box.

Upon connecting the pathway clusters, we uncovered that most reverted genes were linked to lipid metabolism (Node N8) and biological oxidations (N21) (**Supplemental Fig. 4**). These genes had lower expression levels in CD males and ER-agonist-treated males compared to HFD males, consistent with our previous findings (**Fig. 2** and **Supplemental Fig. 2**). While most pathways showed similar effects with both ERα and ERβ activation, we noticed that lipid metabolism was slightly more changed by ERα. Within lipid metabolism, ERα particularly modulated fatty acyl-coenzyme A biosynthesis processes (**Supplemental Fig. 3**). We also uncovered ERβ-dominant effects in regulating phagocytosis (N4), extracellular matrix (ECM, N5), carbohydrate metabolism (N6) and G protein-coupled receptor signaling (N7) (**Fig. 3A**). ERβ agonists specifically suppressed insulin-like growth factor regulation and ECM-related processes such as collagen formation (**Supplemental Fig. 3** and **4**).

Altogether, our analysis revealed extensive implications of systemic ER activation on central processes beyond lipid metabolism and enabled us to distinguish between shared and ER isoform-specific regulation.

### HFD and ER activation signatures co-occur in the liver and are maintained between mouse and primates

Physiological functions of the liver rely on coordinated actions between different cell types. To determine which cell types were affected by HFD and recovered upon the ER-agonist treatments, we analyzed single-cell (comprising 483,955 cells) and spatial transcriptomics datasets [27].

After filtering for males, we focused on cells representing 16 annotated cell types (**Supplemental Fig. 5A**). HFD led to a reduction of major liver cell types, including hepatocytes, endothelial and Kupffer cells, while immune cell populations increased (**Supplemental Fig. 5B**). This confirms previous findings and partly explains HFD-induced gene expression changes in the liver [27,42]. By examining cell type-specific gene expression patterns of our HFD and ER-agonist treatment-derived signatures, we found that the non-reverted, reverted and ERα-specific gene sets (**Fig. 2B**) were mainly enriched in hepatocytes. In contrast, ERβ-specific effects were prominent in endothelial and stromal cell populations, aligning with the profound effects of ERβ on ECM-related genes, including many collagen genes (**Fig. 3B**). These observations potentially reflect different ERα and ERβ activities in hepatic cell populations [43]. The same cell types were affected when mapping these gene signatures to reference macaque and human single-cell liver atlases. This suggested that hepatic molecular key signatures and cellular architecture altered by HFD or in NAFLD are partially maintained between mice and primates, and gene regulatory responses to estrogen treatment are similar in humans (**Supplemental Fig. 5C**).

Analyzing spatial transcriptomics data allowed to identify zonation-specific expression patterns of these signatures across the liver lobule, we found that HFD-induced changes were concentrated near the central vein area, while ERβ-specific effects were enriched in the vasculature including capsule, portal, and central vein (**Fig. 3C**).

To characterize the biological roles of individual cell types in the liver, we assessed the enrichment of previously altered pathways (**Fig. 3A**). We observed that metabolic and oxidative processes occurred in pericentrally located hepatocytes, while processes related to extracellular matrix remodeling operated in stromal cells and the vasculature (**Fig. 3D** and **Supplemental Fig. 5, D and E**). Additionally, comparing pathway enrichment scores from the control to the HFD condition revealed gene expression changes in immune cells promoting phagocytosis and complement cascade processes (**Fig. 3D**).

Overall, our findings highlight that HFD primarily perturbed hepatocyte homeostasis by altering crucial metabolic and oxidative processes, leading to mobilization and activation of immune cells. We find that these gene signatures are maintained in mouse, macaque and human, and that systemic ER activation protects the liver by counteracting these changes.

### Activation of ER-responsive pathways is mediated through changes in chromatin accessibility

The epigenomic and transcriptomic landscapes are intricately linked to maintain cellular homeostasis and can be altered by dietary changes [44]. To investigate ER-agonist-dependent epigenomic restoration of physiological and transcriptional profiles, we performed ChIP-seq on livers of untreated as well as E2 (ERα) and DPN (ERβ)-treated HFD male mice. We focused on modified histones associated with accessible chromatin at promoters (H3K27ac and H3K4me3) and enhancers (H3K27ac and H3K4 monomethylation, H3K4me1). We identified 13,220 promoters and 28,392 enhancers, of which 182 promoters and 1,816 enhancers were differentially acetylated (DAc) at H3K27 upon HFD (**Supplemental Fig. 6, A and B**). Most of these sites gained H3K27ac in response to HFD (57% at promoters and 67% at enhancers), which were partly restored upon ERα and ERβ treatments (**Fig. 4A** and **Supplemental Fig. 6C**).

Enhancer-promoter interactions through chromatin loops impact gene transcription [45], therefore we examined the involvement of DAc enhancers in regulating nearby HFD-affected genes. Overall, we identified 5,448 differentially regulated enhancer-gene (E-G) pairs, of which 68 were estrogen-sensitive (ES-E-G) with 45 unique paired genes residing within chromatin loops (**Supplemental Fig. 6D** and **Supplemental Table 8**). These 45 genes were significantly enriched in metabolic processes (**Supplemental Fig. 6E** and **Supplemental Table 5**), aligning with the observed transcriptomic changes (**Supplemental Fig. 2**).

Among the 68 ES-E-Gs, four enhancers near the *TEA domain transcription factor 1* (*Tead1*) gene showed HFD-induced gain of H3K27ac, which was reduced upon estrogenic ligand treatment (**Fig. 4B**). Using promoter capture Hi-C (CHi-C) data, we discovered interactions between *Tead1* and nearby HFD-regulated gene loci through enhancers and chromatin loop formation (**Fig. 4B**). Additionally, we found enhancers across the Acyl-CoA thioesterase (*Acot*) gene loci that were topologically connected via chromatin loops involving the HEAT Repeat Containing 4 (*Heatr4*) gene locus (**Supplemental Fig. 6F**), suggesting a shared regulatory module for several *Acot* genes.

Combined, these results showed that HFD induces major epigenomic rearrangements in livers of male mice and identified 68 ES-E-Gs that provide insights into the regulatory mechanisms involved. Importantly, these alterations are reversible by ER activation, providing a promising basis for therapeutic interventions.

### Expression of ES-E-G genes in humans is indicative of NAFLD and fibrosis stage

Recent liver cohort studies were designed to identify potential biomarkers and drivers of NAFLD. We reanalyzed a large NAFLD cohort dataset [36] (n=216) to examine the expression levels of ER-reverted orthologs (38/45 genes) in NAFLD patients separated by disease severity (CTRL, NAFL and NASH, collectively termed NAFLD stages) (**Fig. 5A** and **Supplemental Table 9**). By applying *k*-means clustering (*k*=4) to the gene expression profiles, we identified gene sets that were upregulated (cluster 1, n=19) or downregulated (cluster 2, n=7) with disease progression, as well as genes primarily induced in NASH (cluster 3, n=3) or NAFL (cluster 4, n=9) (**Fig. 5A**, **panel 1**). These gene expression patterns correlated with the NAFLD activity score (NAS) spectrum (**Fig. 5A**, **panel 2**) and exhibited limited consistency across fibrosis stages (**Fig. 5A, panel 3**).

**Fig. 5.**
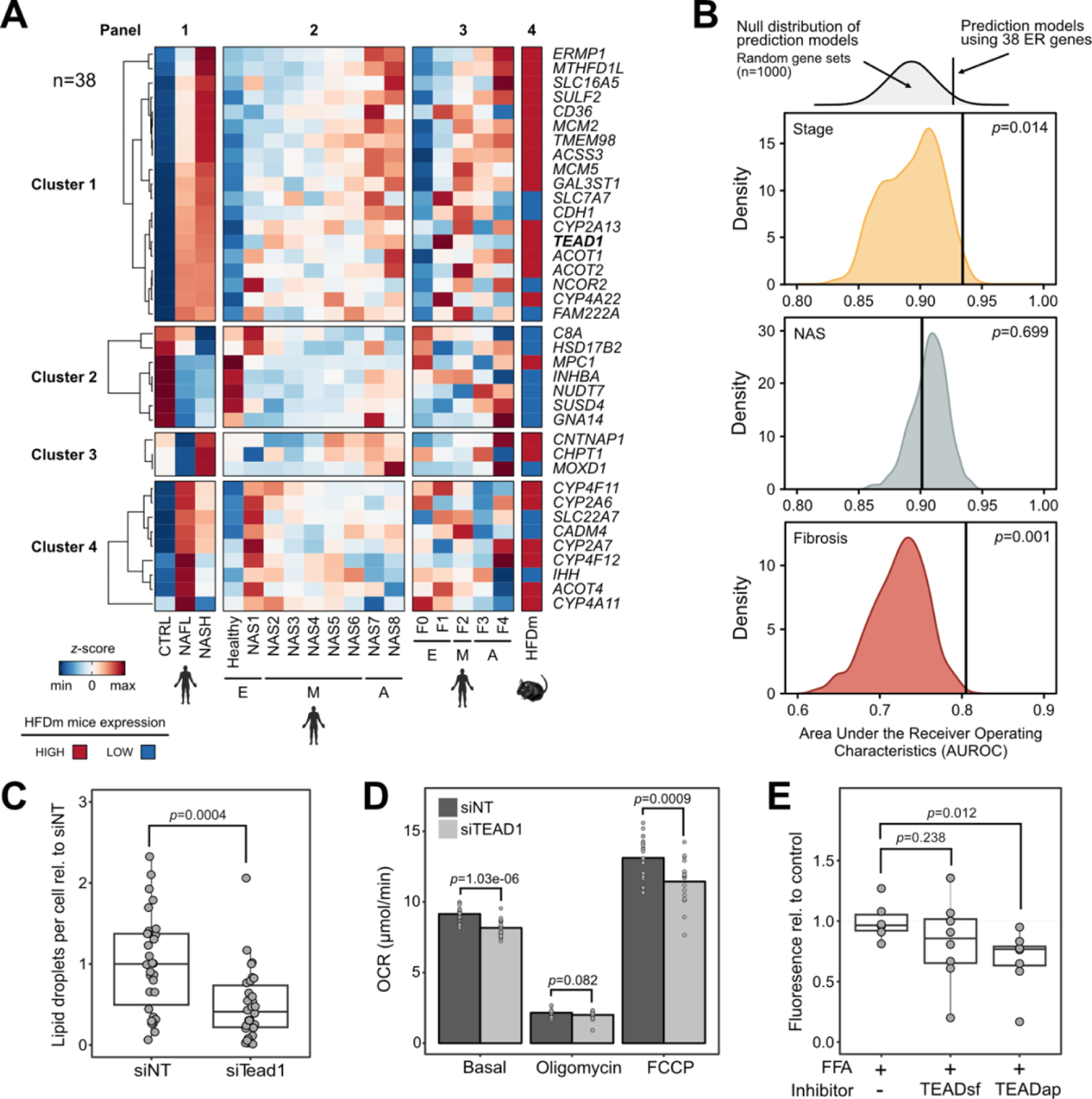
ER-sensitive genes are associated with NAFLD progression and reveal TEAD1 as a clinical target. **(A)** Heatmap displays changes in expression levels for the 38 orthologous ES-E-G genes in NAFLD patients (panel 1-3) and mice (panel 4). Color gradient indicates *z*-score-normalized gene expression counts (blue: low and red: high). Four *k*-means clusters group genes by expression in healthy (CTRL), NAFL and NASH patients (panel 1) as well as patients with different NAS (early (E): NAS0-1, moderate (M): NAS2-6, advanced (A): NAS7-8, panel 2) and fibrosis stages (E: F0-1, M: F2, A: F3-4, panel 3). Expression levels of the 38 genes in HFDm are shown (panel 4). Color code distinguishes downregulated (blue) and upregulated (red) genes in HFDm *versus* CDm. Gene names follow human nomenclature. **(B)** Schematic illustration and density plots show the null distribution of areas under the receiver operating characteristic (AUROC) curve obtained from training 1000 models on random gene set permutations (n=38) predicting NAFLD stage (top), NAS (middle) and fibrosis (bottom) score. AUROC value (black vertical line) of the 38 ER-sensitive genes and *p* values are displayed (permutation tests). **(C)** Box plot shows microscopically quantified lipid droplet number in AML12 cells with siRNA-mediated *Tead1*-knockdown (siTead1) relative to control (siNT). Each box indicates interquartile range (IQR), median (horizontal line) and 1.5×IQR (whiskers). *p* value is shown (two-sided t-test). **(D)** Bar plot displays basal, minimum, and maximum oxygen consumption rate in HepG2 cells with control (siNT) and siRNA-mediated *TEAD1*-KD (siTEAD1). Dots indicate single wells across two biological replicates. Significant *p* values are indicated (two-sided t-test). **(E)** Box plot depicts fluorescently measured lipid content in free fatty acid-fed (FFA+) human primary hepatocyte spheroids. Dots indicate individual spheroids. *p* values are shown (two-sided t-test).

To examine the predictive potential of the 38 ER-regulated genes, we trained models on gene expression signatures based on NAFLD, NAS, and fibrosis stage (**Fig. 5B**). By comparing these 38 genes to size-matched random gene sets, we found that ER-regulated genes were significantly more predictive of NAFLD stage and fibrosis (**Fig. 5B**). Notably, there was strong consistency in the directionality of ER-regulated gene expression changes between NAFLD patient and HFD male mice (**Fig. 5A**, **panel 4**).

*TEAD1* gene expression was increased in NAFLD patients and HFD male mice (**Fig. 5A** and **Supplemental Fig. 7A and B**). Unlike the other three gene family members, *TEAD1* is broadly expressed in liver (**Supplemental Table 10**). *TEAD1* encodes a key transcriptional effector of the Hippo pathway, and this pathway has been recently described to regulate liver homeostasis and metabolism [46,47]. ER-agonist treatment in HFD male mice decreased *Tead1* gene expression (**Supplemental Fig. 7C** and **Supplemental Table 10**). siRNA-mediated knockdown of *Tead1/TEAD1* reduced lipid droplets and oxygen consumption rates in cell lines, suggesting changes in energy metabolism (**Fig. 5, C and D**, **Supplemental Fig. 7C** and **Supplemental Tables 11 and 12**). In a physiologically relevant human model, we treated primary human hepatocyte (PHH) spheroid cultures [12] in steatogenic media with TEADap (VT-104), an inhibitor of TEAD auto-palmitoylation disrupting the interaction between TEAD and its cofactor YAP [13] as well as TEADsf (Ex.174), a small molecule inhibitor binding directly to the TEAD surface blocking the YAP/TEAD interface [14] (**Supplemental Table 10**). Notably, we observed a significant reduction in lipid accumulation with TEADap, exhibiting stronger effects than TEADsf (**Fig. 5E**).

Overall, we identified networks of ER-controlled genes that overlapped between mouse and human livers and were predictive of NAFLD and fibrosis stages. Among the ER-targets genes that showed similar responses was *Tead1/TEAD1*. In an organotypic human liver model, TEAD inhibition reduced hepatic steatosis.

### Hepatic TEAD inhibition ameliorates steatosis by altering central metabolic pathways

To investigate the molecular changes underlying the reduction of hepatic steatosis by TEAD inhibition, we determined gene expression changes in PHH spheroids treated with the TEAD inhibitors in steatogenic media (**Supplemental Table 13**). The TEADap inhibitor induced more DEGs (n=435) compared to the TEADsf inhibitor (n=175), with 125 DEGs shared between both treatments (**Fig. 6A**), indicating that both compounds affected similar genes, albeit to different degrees. DEG analysis revealed a large set of repressed genes (cluster 1, n=391) and a smaller set of activated genes (cluster 2, n=94) (**Fig. 6B**). Pathway analysis (KEGG) of TEADap deregulated genes revealed alterations in molecular metabolism, including AMP-activated protein kinase (AMPK) and phosphatidylinositol-3-kinase (PI3K)-AKT signaling (**Fig. 6C, Supplemental Fig. 8, A and B,** and **Supplemental Table 5**), overall resembling a starvation response. TEAD inhibition may disrupt the direct binding of TEAD proteins to promoters of metabolic genes, for example, *SREBF1* (de novo lipogenesis), *HMGCR* (cholesterol synthesis), or *GHR* (growth hormone receptor), and thereby alter cellular energy and lipid homeostasis (**Fig. 6C** and **Supplemental Fig. 8C**).

**Fig. 6.**
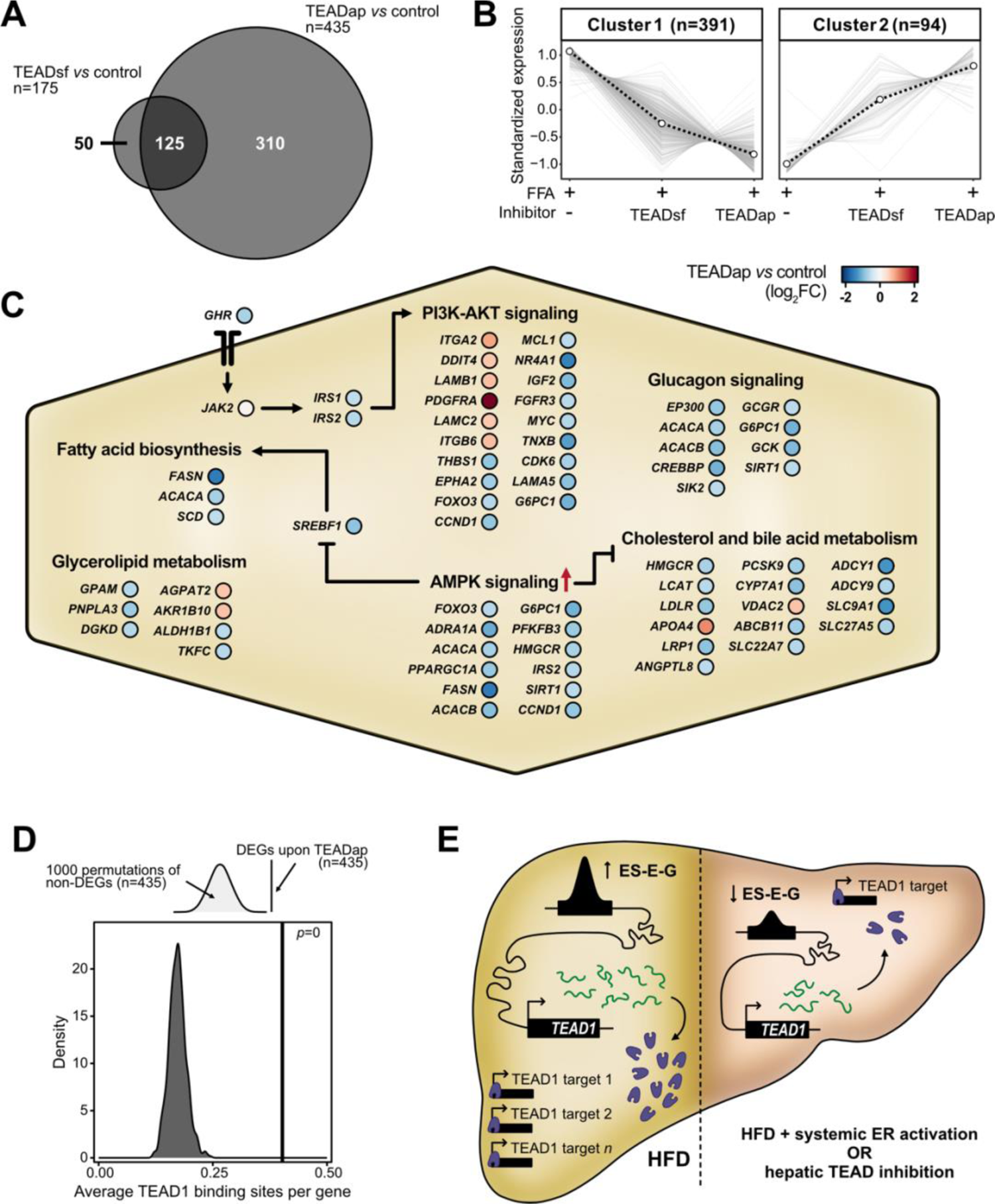
TEAD1 controls core metabolic processes and lipid accumulation in the liver. **(A)** Two-way Venn diagram intersects number (n) of DEGs upon TEADap and TEADsf inhibitor treatments compared to control free fatty acid-fed (FFA+) spheroids. **(B)** Line charts illustrate two clusters of *z*-score-scaled expression trends of unified DEGs (n=485) in FFA+ spheroids without and with TEAD inhibitor treatments. Dashed black line indicates cluster centroid over all deregulated genes (grey). Number indicates genes per cluster (parentheses). **(C)** Schematic illustration displays core pathways altered by TEAD inhibition in FFA+ spheroids. Circles show individual genes in TEADap-treated compared to the untreated condition (blue: reduced, red: increased log_2_FC). Red arrow indicates TEADap-mediated activation of AMPK signaling. Sharp and blunt arrows show activation and inhibition, respectively. **(D)** Schematic illustration (top) and density plot (bottom) show average TEAD1 binding site distributions in promoters of 1000 random non-deregulated gene sets (n=435). The average number of TEAD1 binding sites in promoters of deregulated genes upon TEAD1 inhibition (black line) and *p* value are displayed (permutation tests). **(E)** Model illustrates systemic ER activation effects on liver lipid accumulation upon *TEAD1* gene expression. Increased lipid levels promote *TEAD1* gene locus remodeling with higher enhancer activity and *TEAD1* gene expression (left, yellow background). Systemic ER activation with estrogen treatments suppresses *TEAD1* gene locus activity. Consequently, reduced TEAD1 levels can rewire liver metabolism and decrease lipid accumulation. This effect can be recapitulated by impairing TEAD1 activity using small molecule inhibitors (right, red background).

We then assessed the impact of TEAD on gene regulation through chromatin interactions by quantifying TEAD1 binding sites in DEGs. TEADap DEGs had significantly more TEAD1 binding sites (mean=0.4 per gene) compared to random size-matched gene sets (mean=0.17 per gene, range: 0.12-0.25) (**Fig. 6D**), suggesting direct regulation by TEAD1 rather than secondary signaling mechanisms.

Lastly, we evaluated the contributions of TEAD inhibition to the observed beneficial effects upon ER-agonist treatment. After ortholog conversion, we found that 27.4% (17/62) of significantly enriched genes in the top KEGG pathways after TEADap treatment of PHH spheroids were also differentially expressed upon ER-agonist treatment in male mice on HFD, compared to random size-matched gene sets (median: 9.7%) (**Supplemental Fig. 8D**). Moreover, the gene expression trends in ER-agonist-treated HFD male mice closely resembled those in TEADap-treated PHH spheroids (**Supplemental Fig. 8B**), suggesting that ER-agonist treatment partially restored NAFLD in a TEAD-dependent manner.

To summarize, we demonstrated that systemic estrogen signaling suppresses *Tead1* gene expression in HFD male mice and inhibition of TEAD reduces lipid accumulation in human hepatocytes by repressing crucial lipogenic pathways (**Fig. 6E**).

## DISCUSSION

Beyond reproductive roles, estrogen signaling also maintains tissue homeostasis and estrogenic benefits are well recognized in postmenopausal women and men [3,48]. Similarly, estrogen treatment in male mice alleviated metabolic syndrome, including steatosis and insulin resistance [49]. Our study revealed that estrogenic agonist treatment restored deregulated lipid metabolism and oxidative processes, highlighting the positive metabolic effects of ER activation. Furthermore, we uncovered previously overlooked cellular pathways affected by estrogen signaling, emphasizing its role in maintaining liver homeostasis besides lipid metabolism.

Previous studies involving Estrogen receptor alpha (*Esr1*) gene deletions in both sexes have established ERα as a hepatic key regulator of lipid metabolism, gluconeogenesis and other essential metabolic processes [4,5]. Various dietary disease models have confirmed the protective role of ERα in NAFLD. However, the functional impact of hepatic ERα in safeguarding the liver upon dietary stress remains ambiguous, reporting its requirement [7,8,49,50] and dispensability [50–52]. This ambiguity may stem from developmental shifts in metabolic regulation that affect adult liver function. In our study, ER-agonist treatments in adult mice eliminated congenital confounders and revealed that ERβ activation overall mirrors the cellular and molecular phenotypes observed for ERα signaling [53]. While ERβ is not expressed in hepatocytes (**Supplemental Table 14**), it likely regulates hepatic metabolism through other cell types, such as immune cells present in the liver [42]. Estrogens possess anti-inflammatory properties [54], suggesting that ERα and ERβ contribute to liver homeostasis through immune cells or systemic anti-inflammatory signaling pathways. Our single-cell analysis confirmed that HFD induces inflammatory signaling and alters hepatic immune cell composition, potentially amplifying the responsiveness to or effects by estrogens due to increased proportions of immune cells [42]. Notably, ER-agonist treatments restored the expression of genes involved in monocyte recruitment and inflammatory signaling in HFD male mice.

Low ERβ gene expression was detected in hepatic stellate cells (HSCs) [43], which contribute to fibrosis upon activation. ERβ could mitigate HSC activation and attenuate liver fibrosis. Although our HFD model did not induce fibrosis, the ERβ agonists specifically and predominantly suppressed a range of genes associated with the extracellular matrix, angiogenesis and growth factor signaling. Many of these genes are known to be markedly upregulated upon HSC activation during fibrosis, including Collagen Type I and Type III Alpha 1 Chain (*Col1a1* and *Col3a1)* [55]. Treatment with ERβ agonists may pose a future treatment strategy for diet-induced fibrosis. The transmembrane-bound G protein-coupled ER (GPER1), which can be activated by E2 and PPT but not by DPN, may partially mediate the effects observed with E2 and PPT [5]. Although GPER1 expression was undetectable in our mouse liver data, previous reports demonstrated that GPER1 deficiency in male mice leads to dyslipidemia [56]. Future research involving cell-specific deletions of ERα, ERβ and GPER1 will be needed to dissect the crosstalk between different cell populations and tissues.

Our study revealed that systemic ERα and ERβ activation reversed HFD-induced alterations in enhancer and promoter accessibility. Identifying enhancers is clinically relevant since they can be targeted therapeutically by interfering enhancer RNA [57,58]. We determined a stringent set of 68 ES-E-Gs, including genes associated with NAFLD. Notably, we discovered a large enhancer locus near the *Acot* genes with multiple ERα binding sites, suggesting direct regulation by ERα in the liver. These genes regulate β-oxidation through peroxisome proliferator activated receptor alpha (PPARα) [59], thus linking ERα activation to lipid catabolism. Furthermore, we found four enhancers near the *Tead1* gene locus, which exhibited increased activity by HFD and restoration upon estrogenic ligand treatment. In NAFLD, the Hippo pathway co-factors YAP and TAZ have recently been investigated [46,47], however, the roles and regulation of TEAD1 has been largelyunexplored, mainly due to embryonic lethality upon knockout [60]. Moreover, the involvement of Hippo pathway in energy metabolism in cancer cells promoted the development of drugs targeting oncogenic TEAD which could be repurposed for NAFLD [60].

Many NAFLD treatments targeting metabolic regulators showed efficacy in mice but failed in clinical trials [61]. Therefore, our study focused on identifying genes regulated similarly in mouse and human. Most of our mouse gene candidates exhibited consistent gene expression trends in human, suggesting translatable responsiveness to estrogenic ligand treatment. Specifically, *TEAD1* exhibited similar gene expression trends in HFD male mice and human NAFLD patients. The Hippo pathway, known for regulating tissue homeostasis, is implicated in metabolic disease [46]. Our findings support the notion that Hippo signaling, through TEAD deregulation, activates catabolic metabolic pathways, including cholesterol and fatty acid synthesis upon energy surplus. Inhibiting TEAD and its interaction with YAP presents a promising new therapeutic strategy for metabolic diseases, like NAFLD, bypassing potential adverse effects of estrogen treatment.

## CONCLUSIONS

By integrating multiple datasets in a pre-clinical model, we uncovered that systemic ERα or ERβ activation in HFD male mice remodels the hepatic transcriptome and epigenome, reinstating cellular homeostasis comparable to male mice fed a CD. HFD and systemic ER activation altered core liver pathways, beyond lipid metabolism, that are consistent between mice and primates. Estrogen-sensitive enhancers regulate central metabolic genes of clinical significance in NAFLD patients, identifying *TEAD1* as key ER-sensitive gene and its downregulation by short interfering RNA reduced intracellular lipid content. Subsequent TEAD small molecule inhibition improved steatosis in primary human hepatocyte spheroids by suppressing lipogenic pathways. Thus, the inhibition of TEAD1 has emerged as a new therapeutic candidate for reducing hepatic steatosis while bypassing the pleiotropic effects of estrogen signaling.

## METHODS

### Animal experiments and tissue preparation

Animal experimentation has been previously reported [11]. In short, five-to six-week-old male and female C57BL/6J mice obtained from in-house breeding were fed a control (D12450J, 10% kcal fat, Research Diet) or high-fat diet (D12492, 60% kcal fat, Research Diet) *ad libitum* for 13 weeks. Subsets of male and female mice on HFD were additionally injected intraperitoneally with the estrogenic ligands 17β-estradiol (E2, 0.5mg/kg body weight, Sigma-Aldrich), 4,4’,4’’-(4-Propyl-[1*H*]-pyrazole-1,3,5-triyl)*tris*phenol (PPT, 2.5mg/kg body weight, Tocris), 2,3-Bis(4-hydroxyphenyl)propionitrile (DPN, 5 mg/kg body weight, Tocris) and 4-(2-(3,5-dimethylisoxazol-4-yl)-1H-indol-3-yl)phenol (DIP, 10mg/kg body weight) or given a sham injection every second day from week 10 to week 13. The ligands were diluted in 55% water, 40% PEG400 and 5% DMSO. Mice in each group were descended from different parents and were housed in at least two different cages. Upon sacrifice, livers of C57Bl/6J mice were dissected and washed with phosphate-buffered saline (PBS). Livers were either cross-linked for ChIP-seq, embedded for histology or flash-frozen in liquid nitrogen for RNA-seq. Blood glucose was measured after 2h fasting with a glucometer (Accu-Chek).

### Liver histology of murine liver sections

Formalin-fixed and paraffin-embedded livers were processed into 3µm thick sections, before staining with Hematoxylin & Eosin (Mayers Hematoxylin Plus #01825 and Eosin ready-made 0.2% solution #01650) according to standard histological procedures for the assessment of the liver histology.

### Cell culture

HepG2 and AML12 cell lines were obtained from the American Type Culture Collection with certified genotype and were regularly tested for mycoplasma (Eurofins Genomics). HepG2 cells were cultured in Dulbecco’s Modified Eagle Medium (DMEM) supplemented with 10% fetal bovine serum (FBS, Hyclone, GE healthcare) and 1% Penicillin-Streptomycin (PS, Sigma-Aldrich) while AML12 cells in DMEM/F-12 (Gibco) supplemented with 10% FBS, 1% PS, 1% Insulin-Transferrin-Selenium Sodium Pyruvate (Gibco) and 20ng/mL Dexamethasone (Sigma-Aldrich) in T75 flasks at 37°C and 5% CO_2_ atmosphere. Cells were passaged at a 1:6 ratio twice (HepG2) and three (AML12) times a week by aspirating the medium, gently washing the cells with PBS without Mg^2+^ (Sigma-Aldrich) and then detached using 2mL of trypsin-EDTA solution (Sigma-Aldrich) for 3-5min. Trypsin was inactivated with 8-10mL of culture medium before passaging to a new flask.

### Primary human hepatocyte spheroid culturing

Spheroids were formed by seeding cryopreserved primary human hepatocytes (PHH) of a male donor in ultra-low attachment 96-well plates (Corning) as previously described [12]. For spheroid treatments, free fatty acids were conjugated to 10% bovine serum albumin at a molar 1:5 ratio for 2h at 40°C. Formed spheroids were treated with 240μM oleic acid and 240μM palmitic acid along with 100nM of either TEAD autopalmitoylation (TEADap, VT-104, 100nM) [13] or TEAD surface inhibitor (TEADsf, 100nM) [14] inhibitors for 5 days. Intracellular lipid content was assessed using the AdipoRed Assay Reagent (Lonza).

### siRNA-mediated TEAD1/Tead1 knockdown

Confluent HepG2 (60-70%) or AML12 (80-90%) cells were trypsinized and electroporated with siRNAs targeting either TEAD1/Tead1 (SMARTpool) or a control non-targeting siRNA pool (ON-TARGETplus^TM^, Horizon Discovery). After washing cells once with OptiMEM (Invitrogen), 2 million cells were resuspended in 200μl OptiMEM and incubated for 3min with 2μg (7.5μL of a 20μM stock) siRNA in a 4mm cuvette (Bio-Rad) before being pulsed at 300V, 250μF, in a Genepulser II (Bio-Rad). Immediately after electroporation, the cells were transferred to pre-heated (37°C) phenol red-free DMEM (HepG2) or DMEM/F-12 (AML12) culture medium without antibiotics. Cells were collected at day four to determine knockdown efficiency and microscopy, and at day five for Seahorse analysis.

### Microscopic LD quantification

Two days after electroporation, 30,000 AML12 cells transfected with siNT or si*Tead1* were seeded into ibiTreat 8-well cover slips (ibidi). The following day, LDs were stained with LipidTOX Red (Thermo Fisher Scientific) 1:6250 (v:v) and nuclei with NucBlue (Thermo Fisher Scientific) 1:62.5 (v:v). After incubation at 37°C and 5% CO_2_ for 20min, the cells were washed twice washed twice with Leibovitz’s L15 medium. Images were acquired using a LSM780 confocal microscope (Zeiss) with a Zeiss C-APOCHROMAT water immersion objective lens (40x /1.2). Imaging was performed at 37°C. NucBlue and LipidTOX Red were excited using 405nm and 640nm laser lines, respectively. Image analysis was carried out using ImageJ. Nuclei were identified and subsequently counted by masking the NucBlue channel after applying a 3-pixel mean filter. Individual LDs were located by identifying local intensity maxima after applying a 2-pixel mean filter in the LD channel. To quantify the average number of LDs per cell, the total number of identified LDs in each image was divided by the nucleus number.

### Seahorse assay

Metabolic flux analysis was carried out on HepG2 cells using Seahorse XF96 Extracellular Flux Analyzer (Agilent). 15,000 cells were seeded the day before the experiment, and medium was changed to XF DMEM based medium containing 2mM GlutaMax, 25mM glucose and 1mM pyruvate on the day of the experiment and incubated at 37°C without CO_2_ for 1h prior to the experiment. The oxygen consumption rate was measured at and following injection of oligomycin (1µM final), FCCP (0.5-1.5µM), and mixture of rotenone and antimycin A (4µM). Data were normalized on the number of cells per well and against basal oxygen consumption rate. Cell number was normalized by nuclear staining (Hoechst, Molecular probes) for 10min followed by imaging each well using BD pathway 855 (BD Biosciences) with a 10x objective and montage 5×4. Cell number was counted with Cell profiler software.

### RNA isolation and DNase treatment

Approximately 20µg of flash-frozen liver tissue was homogenized in 700µL QIAzol (QIAGEN) using a TissueLyzer II (QIAGEN, 2min, 25 Hz, 2 times). The samples were incubated at room temperature for 5min, before adding 140µL chloroform (Sigma-Aldrich). This mixture was shaken for 15s, incubated for 3min, and centrifuged at 9,000×g and 4°C for 5min. The aqueous phase was carefully removed, and an equal volume of isopropanol added. This mixture was incubated at room temperature for 10min, before centrifugation at 20,000×g and 4°C for 10min. Supernatant was removed and the pellet washed twice with 70% ethanol, air-dried, and resuspended in water. The isolated RNA was DNase treated using the Turbo DNase Kit (Thermo Fisher Scientific) according to the manufacturer’s instructions. In brief, 10µg of RNA was treated with 2U DNase and 40U RNaseOUT (Thermo Fisher Scientific) at 37°C for 30min. DNase-treated RNA from mouse livers was incubated with DNase inactivation reagent (Turbo DNase kit) for 5min under constant homogenization. The sample was centrifuged at 10,000×g for 2min to remove the inactivation reagent. To purify the obtained DNase-treated RNA, the RNA was diluted to 130µL with water, before adding 20µL sodium acetate (Thermo Fisher Scientific, 3M, pH 5.2), 1µL GlycoBlue (Thermo Fisher Scientific) and 600µL ice-cold 99.8% ethanol. Next, RNA was precipitated at −80°C overnight, before centrifugation at 20,000×g for 30min, washing the pellet twice with 70% ethanol, air-drying and resuspending in water. DNase-treated RNA from liver spheroids was purified using an RNA clean and concentrator kit according to the manufacturer’s instructions (Zymo research). The RNA quality was assessed on a Bioanalyzer 2100 device using RNA Nano chips (Agilent Technologies) and only high quality RNAs (RIN > 6.5) were used for RNA-seq.

### RNA sequencing and data processing

Strand-specific RNA libraries were generated using the NEBNext Ultra II stranded library kit (New England Biolabs) combined with polyA-coupled beads (New England Biolabs) according to the manufacturer’s instructions. The library quality was assessed on a Bioanalyzer 2100 device using DNA High Sensitivity chips (Agilent Technologies) and quantified using a KAPA library quantification KIT (Roche). cDNA libraries were subsequently sequenced on an Illumina NextSeq 500 device using a paired end high output kit (75+75 cycles for mouse, 40+40 cycles for PHH spheroids). Reads were trimmed (Trimmomatic v0.36 for mouse, fastp v0.23.2 for PHH spheroids) and filtered for non-ribosomal RNA by mapping to a custom rRNA reference (HISAT2 [15] v2.1 for mouse, SortMeRNA v4.3.6 for PHH spheroids). Non-aligned reads were further mapped to the mm10 mouse reference genome retrieved from GENCODE vM23 (HISAT2, GRCm38.p6) or Ensembl release 109 in the case of PHH spheroids (HISAT2 v2.2.1). Generated SAM files were converted to BAM files and consequently processed (SAMtools [16] v1.9 for mouse, SAMtools v1.16.1 for PHH spheroids). bedGraph files were generated using HOMER [17] (v4.10) for mouse or bedtools (v2.30.0) for PHH spheroids. Count tables were generated using SubRead [18] (v1.5.2 for mouse, v2.0.3 for PHH spheroids).

### Differential gene expression analysis

Differential gene expression analysis for mouse RNA-seq data was performed using DESeq2 [19] (v1.30.0, default model) and edgeR [20] (v3.32.1, glmFit model). Genes which were found to be differentially expressed in both analyses were considered for further analysis. Human spheroid RNA-seq data was analyzed with DESeq2 (v1.38.3, default model).

### Transcriptomic signal-to-noise ratio

Transcriptome-wide differences across conditions were measured unbiased by using a transcriptome-based signal-to-noise ratio (tSNR). For this, the Euclidean metric was used as a measure of distance across averaged transcriptomes between groups while the noise was defined based on the total within-group variation observed, expressed as:

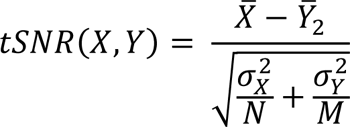

Here, *X^-^* and *Y^-^* are the averaged transcriptomes, *N* and *M* indicate the sample number, and σ^2^_X_ and σ^2^_Y_ represent the intragroup variance for *X* and *Y* groups, respectively.

### Gene clustering and overrepresentation analysis

To identify shared gene expression patterns across the different diet and agonist-treated conditions, we applied a soft clustering strategy using the Mfuzz [21] R package (v2.48.0). Normalized gene expression values were *z*-score scaled (*µ*=0; sd=1) and the optimal number of clusters was determined after testing for different numbers of clusters. Gene ontology (GO) and Kyoto Encyclopedia of Genes and Genomes (KEGG, release 106) analyses on selected gene sets were performed using hypergeometric tests with the hypeR [22] or clusterProfiler [23] R packages. GO biological process annotations were retrieved from the MGI database (v6.16; 03-2021), org.Mm.eg.db (v3.12.0 and v3.16.0) or org.Hs.eg.db (v3.16.0), and enriched terms were established using a *q* value threshold of 0.05 and a custom gene background. The rrvgo [24] package was used to cluster and produce visual representations of the overrepresented terms. Semantic similarities between terms were calculated using the Wang method and a similarity threshold of 0.8 was used for determining GO clusters.

### Pathway enrichment and network analysis

Gene set enrichment analysis (GSEA) was used to identify enriched Reactome pathways between conditions. The fgsea R package (v1.14.0) was run with gene lists ranked according to the signed log_10_ *p*-values obtained from DESeq2 and pathway sizes were limited to a range of 10-500 genes relative to the background. Here, a *q* value threshold of 0.05 was applied for enriched pathway selection. Network visualizations of all enriched pathways were generated using Cytoscape [25] (v3.8.2). Individual pathways were connected according to their similarity (*s* > 0.5) and clusters of the interconnected pathways were produced using the GLay community clustering algorithm from clusterMaker [26]. To uncover shared or divergent trends across different processes, pathway clusters were correlated based on their average normalized enrichment score (NES) and connections were filtered to those with |*r|* > 0.8. The similarity score between pathways used is a metric of both the jaccard similarity and overlap coefficients, calculated as:

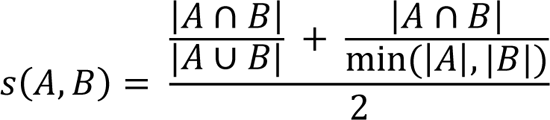

Where *A* and *B* represent the two sets of genes that are part of the pathways being compared.

### Single-cell data analysis

Public single-cell and spatial transcriptomics datasets were retrieved from the Liver Cell Atlas [27]. Only male mouse samples were used for the single-cell reference map and primary cells were removed. Furthermore, only data from male macaque and human were considered. Cell type composition analyses were conducted in R using Seurat [28] (v4.0.2). Enrichment scores for the relevant ER activation signature gene sets and Reactome pathway clusters identified were calculated using pagoda2 (v1.0.2). Scores were aggregated at the cell type level by averaging the enrichment values of all individual cells annotated for a given cell type cluster and condition.

### ChIP-sequencing and data analysis

Formaldehyde-fixed livers were homogenized using a douncer and washed twice with ice-cold PBS. Nuclei were prepared as previously described [29] and sonicated using a Sonics Vibra cell VCX 750 set to 32% duty cycle for 30 cycles (30s on, 59s off). ChIP was performed using antibodies against H3K27ac (Abcam #4729, lot: GR3211959-1, 5µg) and H3K4me3 (monoclonal, Merck 05-1339, lot: 3305923, 5µg) as previously described [29]. Libraries from immunoprecipitated DNA were generated using the SMARTer ThruPLEX DNA-seq Kit (Takara Bio), size-selected and quality assessed by Bioanalyzer DNA High Sensitivity chips (Agilent Technologies) according to manufacturer’s protocols. Libraries were quantified using KAPA quantification kit (Roche) and sequenced on an Illumina NextSeq 500 device using a single end (75 cycles) high output kit. Reads were mapped to the mouse reference genome (GRCm38.p6/mm10, bowtie2 [30] v2.3.5.1), processed and sorted (SAMtools v1.12), masked regions (NGSUtils v0.5.9) and duplicate reads were removed and indexed (SAMtools v1.12). Peaks were identified using MACS [31] (v2.2.6). bedGraph files were generated (deepTools [32] v3.3.2) and differentially bound peaks were determined (DiffBind v3.0.15) with the threshold FDR < 0.05 and |log_2_FC| > 0.585. Selected regions were annotated (ChIPpeakAnno [33] v3.24.2). Raw H3K4me1 (E-MTAB-7127), CTCF (E-MTAB-437) and Erα (GSE49993) ChIP-seq data were retrieved and processed.

### Quantification of H3K27ac signals in differentially acetylated regions

BED file containing differentially acetylated promoters and enhancers (±200bp from the peak center) was converted into SAF format. H3K27ac BAM files and SAF annotation file were used to generate a count table normalized by counts per million (SubRead v2.0.0).

### Enhancer-gene pair analysis

The closest transcription start sites (one upstream, two downstream) to each differentially acetylated enhancer were determined using BEDOPS [[34] closest-features (v2.4.39). H3K27ac and gene expression (TPM) of single replicates were correlated using Pearson correlation. Only enhancer-gene pairs with H3K27ac to gene expression correlation of |*r*| > 0.75 were considered. Enhancer-gene pairs containing genes recovered by estrogenic ligand treatments were further analyzed. CTCF motif orientation (MA0139.1) in the mouse genome (mm10) was determined using FIMO [35] (MEME Suite). To identify potential canonical or non-canonical CTCF-mediated chromatin loops, we filtered for enhancers harboring an upstream CTCF peak (plus-strand oriented motif for canonical and minus-strand oriented motif for non-canonical loops, within 50kb) when the paired genes were located downstream, and for downstream CTCF peaks (minus-strand oriented motif for canonical or plus-strand oriented motif for non-canonical loops, 50kb) when the paired genes were located upstream of the enhancer. Promoter-capture Hi-C data (GSE155153, Zeitgeber time 0) were lifted over to mm10 to generate contact maps (UCSC LiftOver).

### Patient classification models

Human orthologs of the genes part of estrogen-sensitive enhancer-gene pairs in mice (n=45) were determined using Ensembl release 105. The NAFLD cohort data [36] (gender-balanced) was retrieved from Gene Expression Omnibus (GSE135251) and normalized to counts per million (CPM), scaled and centered (*z*-score). Only human orthologs with gene expression CPM>0.5 were considered (n=38). Multiclass classification models were trained using the glmnet method from the caret R package [37] (v6.0-92) and a split of 3:2 was used for obtaining datasets for training and testing. For NAS and fibrosis categorization, classes were defined as early (NAS0-1, F0-1), mid (NAS2-6, F2) and advanced (NAS7-8, F3-4). Model evaluation was conducted with the multiROC R package (v1.1.1) using the micro average metric for comparison of receiver operating characteristic (ROC) curves across models. Permutation tests were conducted using a null distribution of 1000 models trained by random sampling of 38 genes from the gene background.

### Transcription factor motif search

Genome-wide transcription factor binding sites (TFBS) for TEAD1 were identified using PWMscan [38]. The mononucleotide position weight matrix for the TEAD1 binding motif was retrieved from the HOCOMOCO v11 database [39] for establishing TFBS a background nucleotide composition (A=0.29, C=0.21, G=0.21, T=0.29) and *p* value cutoff of 0.00001 were set. Only TFBS in gene promoters were considered, defined as those within 1.5kb upstream and 0.5kb downstream of transcription start sites. The promoter region with the highest number of TFBS was assigned to each gene.

### Use of standardized official symbols

We use HUGO (Human Genome Organization) Gene Nomenclature Committee-approved official symbols (or root symbols) for genes and gene products, all of which are described at www.genenames.org. Gene symbols are italicized, whereas symbols for gene products are not italicized.

### Statistics

All analyses were conducted in R (4.0 or 4.2). The Shapiro-Wilk test was used to assess normality of the data.

## DECLARATIONS

### Ethics approval and consent to participate

All experimental protocols were approved (N230/15) by the local ethical committee of the Swedish National Board of Animal Research.

### Authors’ contributions

MKA, HH, VML, AA, CW and CK conceptualized the project. CSa, LH, MB, RI, JNS, MKA, AA and CK performed *in vivo* experiments. CSo, JXS, CFA and GM performed *in cellulo* experiments. CSo, CSa, PC, KG, ES and CK carried out tissue and cell stainings or the microscopic imaging. CSo performed RNA-seq and ChIP-seq experiments. CP provided TEAD inhibitors. HH and VML provided human sample material. CSo and CGD analyzed and visualized the mouse and cohort RNA-seq datasets. CSo analyzed and visualized the ChIP-seq data. CGD performed the pathway enrichment and scRNA-seq analyses, and trained machine learning models. MKA, ES, HH, VML, CW, and CK acquired funding and supervised the project. CSo, CGD and CK wrote the original draft. CK acted as guarantor. All authors contributed to the review and editing process.

## Acknowledgements

We would like to thank the group members of the laboratories of Claudia Kutter, Marc Friedländer and Vicent Pelechano for helpful feedback regarding the experimental procedures, data analysis, and data presentation, as well as Noah Moruzzi for assistance with the Seahorse experiment. We thank the Liver Cell Atlas, EPoS and LITMUS consortium for sharing invaluable data with the research community.

## LIST OF ABBREVIATIONS

AUC: area under the curve

CTCF: CCCTC-binding factor

CD: control diet

CHi-C: promoter capture Hi-C

ChIP-seq: chromatin immunoprecipitation followed by sequencing CoA: coenzyme-A

CPM: counts per million DAc: differentially acetylated

DEG: differentially expressed gene

DIP: 4-(2-(3,5-dimethylisoxazol-4-yl)-1H-indol-3-yl)phenol

DPN: diarylpropionitrile

E2: 17β-estradiol

ECM: extracellular matrix

ER: estrogen receptor

ES-E-G: estrogen-sensitive enhancer-gene pair

FCCP: carbonyl-cyanide 4-trifluoromethoxy-phenylhydrazone

GEO: gene expression omnibus

GO: gene ontology

GPCR: G-protein coupled receptor

GSEA: gene set enrichment analysis

H&E: hematoxylin and eosin

HFD: high-fat diet

H3K27ac: histone 3 lysine 27 acetylation

H3K4me1: histone 3 lysine 4 monomethylation

H3K4me3: histone 3 lysine 4 trimethylation

HSC: hepatic stellate cell

KEGG: Kyoto Encyclopedia of Genes and Genomes

NAFLD: nonalcoholic fatty liver disease

NASH: nonalcoholic steatohepatitis

NAS: NAFLD activity score

NES: normalized enrichment score

PCA: principal component analysis

PPT: pyrazole-triol

ROC: receiver operating characteristic

TAZ: WW domain containing transcription regulator 1

TPM: transcripts per million

tSNR: transcriptome-based signal-to-noise ratio

TSS: transcription start site

YAP: Yes-associated protein

